# Evidence that pleiotropic alleles underlie adaptive divergence between natural populations

**DOI:** 10.1101/718916

**Authors:** Ken A. Thompson

## Abstract

The alleles used for adaptation can pleiotropically affect traits under stabilizing selection, and compensatory alleles can be favoured by selection to counteract such deleterious pleiotropy. Such compensatory alleles can segregate in interpopulation hybrids, causing segregation variance for traits where parents have the same phenotype. If adaptation typically involves pleiotropy and compensation, then the segregation variance for traits under stabilizing selection is expected to increase with the magnitude of adaptive phenotypic divergence between parents. This prediction has not been tested empirically, and I gathered data from experimental hybridization studies to evaluate it. I found that pairs of parents which are more phenotypically divergent beget hybrids with more segregation variance in traits where the parents do not differ. This result suggests that adaptive divergence between pairs of natural populations proceeds via pleiotropy and compensation, and that potentially deleterious transgressive segregation variance accumulates systematically as populations diverge.

## Introduction

When populations adapt to their environment, they increase the frequency of (or fix) alleles that affect the phenotypes of traits under selection. The alleles that underlie adaptation can also affect other phenotypes that might be under stabilizing selection, a phenomenon known as pleiotropy (Stearns 2010). In recent years, evidence has accumulated, largely from evolutionary model systems, which suggests that pleiotropy is common (although it might only affect a small subset of organisms’ traits; Wagner et al. 2008; Wagner and Zhang 2011; Wang et al. 2010). If the pleiotropic effects of alleles are deleterious, compensatory mutations that counteract this deleterious pleiotropy can be favoured by natural selection (Phillips 1996). Although this model of adaptation via pleiotropy and compensation emerges in many theoretical models of adaptation (Barton 2001; Orr 2000), it is unclear whether such a process typically characterises adaptation in natural populations.

Predictions from theoretical models of divergent adaptation and hybridization can be tested to infer whether adaptation in natural populations typically involves pleiotropy and compensation. Barton (2001) conducted simulations of Fisher’s (1930) geometric model of adaptation in a case where two populations with ten traits experienced divergent selection on a single trait while while the other nine were subject to stabilizing selection. Following hybridization of the two populations, there was appreciable phenotypic variation in the nine traits under stabilizing selection. This segregation variance was caused by the haploid recombinant hybrids inheriting alternative combinations of compensatory alleles. Importantly, the amount and/or average effect size of compensatory alleles should be positively correlated with the amount of phenotypic divergence between the parents. Thus, the theoretical prediction under adaptation via pleiotropy and compensation is: as the phenotypic divergence between pairs of populations increases, so should the amount of segregation variance in non-divergent traits observed in their hybrids. See Fig. 1 for a visual overview of this prediction and Fig. S1 for the results of computer simulations illustrating the prediction more quantitatively. Here, I test this theoretical prediction using data collated from experimental crossing studies.

**Fig. 1.**
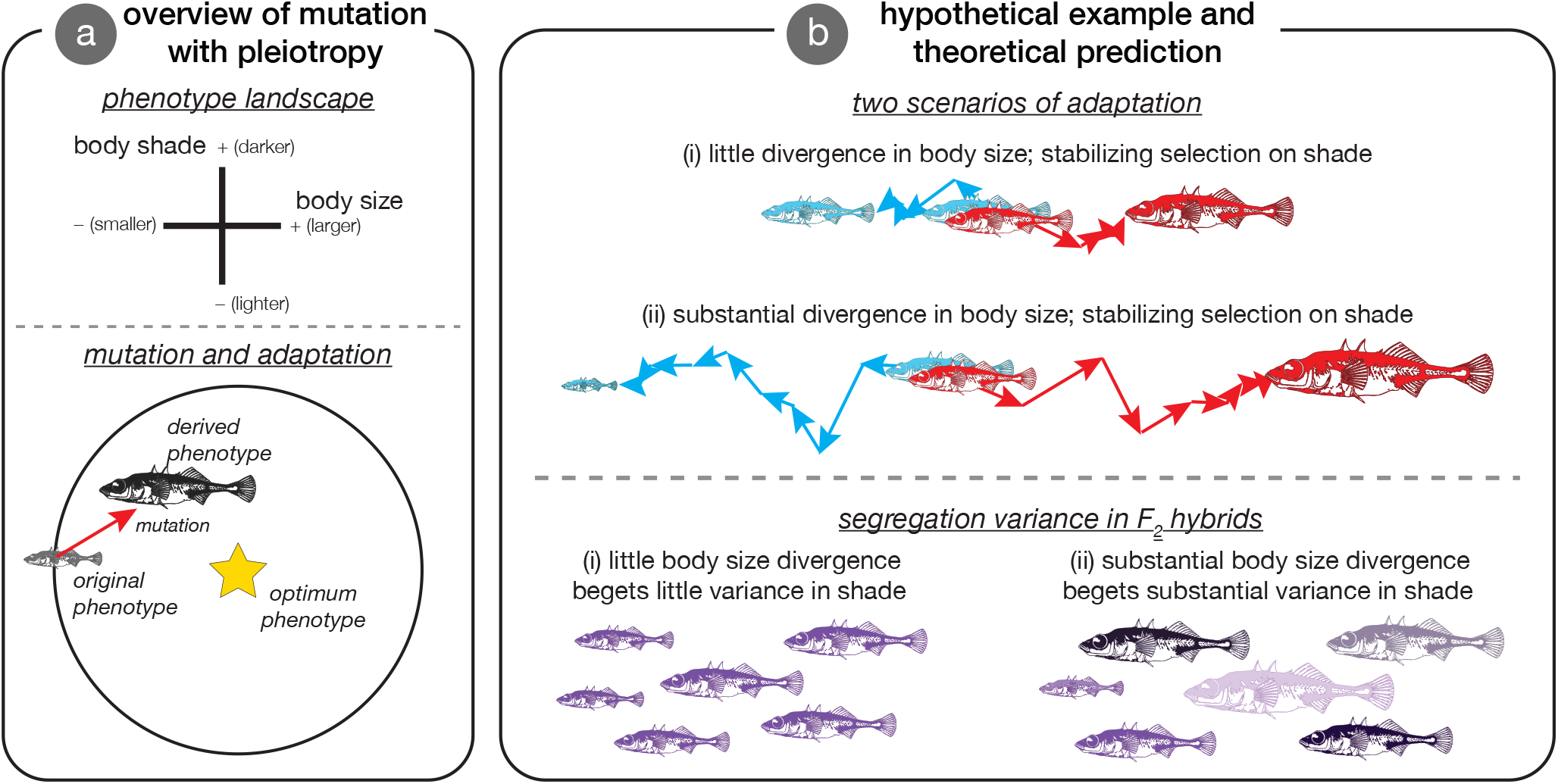
Overview of adaptation with pleiotropic alleles and theoretical prediction. Panel **A** shows a general overview of Fisher’s geometric model, which relies on pleiotropic mutation. The upper section shows the phenotype landscape under consideration, wherein the horizontal axis is body size (small - big) and the vertical axis is body shade (not colour; light - dark). The lower section illustrates the fixation of a pleiotropic allele during adaptation. The original phenotype is medium in size and medium in shade. The optimal phenotype is larger but the same shade. A mutation arises that greatly increases size and has a deleterious pleiotropic effect to darken shade. Since the mutation is beneficial (points inside the circle), it has a high probability of fixation in spite of the deleterious side-effect. Panel **B** illustrates the theoretical prediction in two diverging populations – red and blue – with the same initial phenotype for size and shade—colour here is just used to visually demarcate parent populations and hybrids (purple) and is not considered a trait. The upper section illustrates two alternative adaptive scenarios for comparison. In both scenarios (i) and (ii), shade is under stabilizing selection in the two populations. Scenario (i) is a case where the two populations diverge little in body size and scenario (ii) represents substantial divergence in body size. The lower section of the panel illustrates the outcome of hybridization. The key insight is that the segregation variance in shade is greater in (ii) than (i). Body size segregates as well, but it would do so in a model without pleiotropy whereas shade would not necessarily. Darker recombinant hybrid individuals inherited mostly compensatory alleles that darken shade (i.e., they point ‘up’) and lighter individuals inherited mostly compensatory alleles that darken shade (i.e., they point ‘down’).

## Methods

I conducted a systematic literature search with the goal of identifying studies that measured phenotypic traits and variances in two parent taxa and their F_1_ hybrids in a common environment (as a part of a separate study). To be selected for inclusion, studies had to measure at least one non-fitness trait (i.e., ‘ordinary’ trait [Orr 2001]) and parent taxa had to be fewer than 10 generations removed from the wild (details of the literature search are given in the supplementary methods). In total, I (with help) screened over 11,000 studies and collected data from 198. Of these 198 studies, all that met the following two criteria were included in the present analysis: (1) F_2_ hybrids were measured and (2) the parents had significantly different phenotypes for least one trait and were statistically indistinguishable for at least one other.

After obtaining studies for possible inclusion, I filtered and binned the data to generate summary statistics for analysis. Filtering and binning decisions were ultimately some-what subjective, and I present the test of the main hypothesis for summary datasets generated under alternative filtering and binning criteria in Table S1 — the qualitative conclusions of my analysis are robust to alternative decisions during data processing. In addition, further analysis with potentially low-power studies removed illustrate that the observed patterns are not caused by associations between sample size (number of individuals measured) and any variables (see also Table S1). I also note that methods are briefly (but sufficiently) detailed here in the main text, but full detail with appropriate citations are given in the supplementary methods. All analyses were conducted in R v3.5.1 (R Core Team 2018).

In the main text, I restricted my analysis to morphological traits — by far the most frequently measured trait type in the studies that met the above criteria — to maximize the degree to which traits and units were comparable. In total, I retained 15 crosses from 14 studies for the present analysis (Bradshaw et al. 1998; Bratteler et al. 2006; Hermann et al. 2015; Husemann et al. 2017; Jacquemyn et al. 2012; Koelling and Mauricio 2010; MacNair et al. 1989; McPhail 2008; Mione and Anderson 2017; Pritchard et al. 2013; Raeymaekers et al. 2009; Selz et al. 2014; Shore and Barrett 1990; Vallejo-Marín et al. 2017).

For each study I divided traits into two groups: those that differed between the parents — which I assume was the result of divergent selection — and those that did not and were more likely (though not necessarily) subject to stabilizing selection. I classified traits as divergent if they were significantly different (*P* < 0.05) in a *t*-test. For each trait, I calculated the degree of phenotypic divergence in units of parental phenotypic SDs using the smaller of the two parental values. For each study, I then calculated phenotypic divergence for both groups of traits as the mean of ln-transformed divergence values.

For traits that were statistically indistinguishable, I determined the segregation variance of each as:

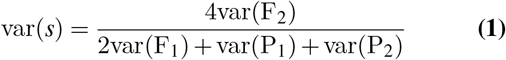

Wright (1968). This quantity normalizes for the standing variation observed in each parent and F_1_ hybrids and captures the variance due to the segregation of population- or species-specific alleles. For each study, I took the mean of these values after ln-transformation as an estimate of segregation variance.

My prediction was that if adaptation commonly proceeds via pleiotropic and compensatory alleles, there should be a positive relationship between parental divergence – for divergently selected traits – and segregation variance – for traits that do not differ between the parents. Visualization of linear models and statistical tests of heteroskedasticity clearly showed that the assumptions of parametric statistical analyses were violated (see Fig. S2). Accordingly, I tested all predictions using Spearman’s rank-order correlations, which test if more divergent pairs of populations beget hybrids with more segregation variance as compared to lesser divergent parental taxa.

A similar pattern to what is predicted above could be the result of genetic drift and have nothing to do with divergent natural selection. Specifically, if more phenotypically divergent populations also have a more ancient divergence time, they might have fixed a greater number of compensatory mutations (if such mutations fix at a steady rate over time). If this was the case, one would detect the predicted pattern even if the alleles underlying divergence were not pleiotropic. It is therefore important to rule out this role for time by testing whether phenotypic divergence is correlated with divergence time in the studies analyzed herein. I did this using three main approaches: (1) by comparing phenotypic divergence of intra-specific c ross p arents v s. i nter-specific cr oss parents, (2) by evaluating the correlation between neutral gene sequence divergence and phenotypic divergence, and (3) by evaluating the correlation between estimates of divergence time and phenotypic divergence.

## Results

I observed a positive correlation between the mean parental phenotypic divergence in statistically divergent traits and segregation variance in statistically indistinguishable traits (Spearman’s *ρ* = 0.800, *P* = 0.000581, *n* = 15) (Figure 2). The phenotypic difference between parents for statistically indistinguishable traits was not correlated with the segregation variance in those traits (Spearman’s *ρ* = 0.446, *P* = 0.0972, *n* = 15) (Figure S3). The patterns were generally robust to data processing decisions (see Table S1), but became non-significant when I included physiological and chemical traits (*P* = 0.052).

**Fig. 2.**
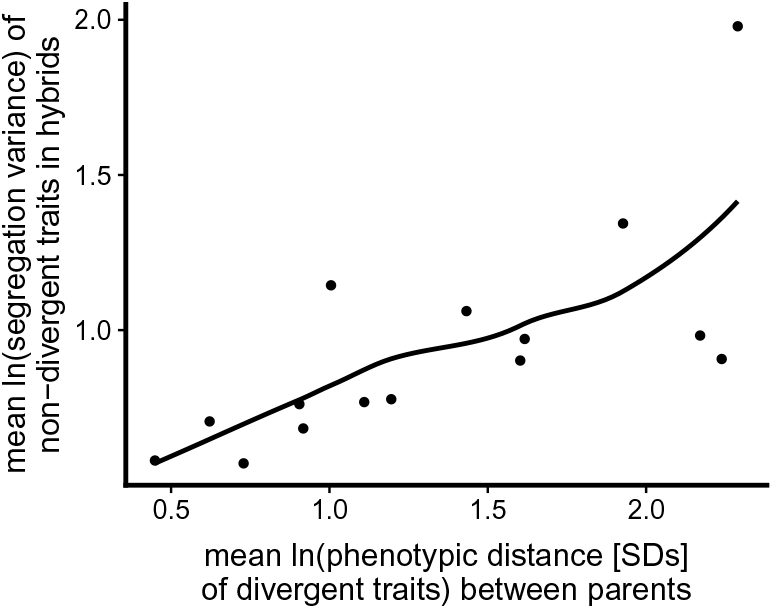
Scatterplot depicting the relationship between phenotypic divergence and transgressive segregation variance for morphological traits. Each point (*n* = 15) represents a unique cross between two populations or species. Points to the right on the horizontal axis have more divergent parents for those traits that are divergent (Spearman’s *ρ* = 0.800, *P* = 0.000581). The line is a loess fit.

Divergence time could correlate with phenotypic divergence between populations, which would render it difficult to disentangle the effects of time and phenotypic divergence. A *t*-test did not detect a difference between intra-specific and inter-specific crosses in phenotypic divergence (F_1,13_ = 0.013, *P* = 0.912) for the studies in our main analysis. Additional analyses found no support for associations between any variable and genetic divergence (Fig. S4C), divergence time (Fig. S4D), or phylogeny (phylogenetic signal test, all *P* > 0.5).

## Discussion

I tested the hypothesis that divergent adaptation is associated with transgressive phenotypic variation in recombinant hybrids. This prediction holds if the genes underlying divergent adaptation are pleiotropic and does not if they are not (or if they are pleiotropic but have infinitely small individual effects (Barton et al. 2017)) (see Fig. S1). Given the lack of effect of divergence time (or its correlates) on phenotypic divergence in the data, the consistency between the results presented here and the theoretical prediction suggests that the genes used during adaptation are indeed pleiotropic and of appreciably large effect. The results might also hint at of the mode of adaptation for the taxa considered herein. For example, adaptation from standing variation causes greater transgressive segregation variance compared to adaptation from *de novo* mutation (Thompson et al. 2019), and thus the observed patterns could be a consequence of adaptive divergence from standing variation. Even if large-effect pleiotropic mutations arise, models with slowly moving fitness optima predict that only alleles with very-small effects will be used during adaptation (Matuszewski et al. 2014). The analyses above suggest that optima in nature move quickly enough for alleles of non-trivial effect sizes to be incorporated.

My findings might initially appear to contradict results of previous empirical studies of transgressive segregation. For example, Stelkens and Seehausen (2009) and Stelkens et al. (2009) found that genetic distance, but not phenotypic distance, predicts transgressive segregation. Although this seems to contradict the pattern shown in Fig. 2, the predictions are not directly comparable I binned traits into categories of divergent & non-divergent and compared parental divergence in the former to hybrid variance in the latter. By contrast, Stelkens’ studies investigated the degree to which individual hybrids are transgressive for traits considered on their own or across all traits. Thus, the analysis presented herein tests a separate hypothesis than has been tested previously.

In Barton’s (2001) simulations, segregation variance in non-divergent traits was deleterious and accordingly hybrid fitness declines as segregation variance increases. It would be useful to test this prediction in an experimental system. Parental lines could be selected for divergence to varying degrees (e.g., differing concentrations of a stressor) and then hybridized with a common ancestor. F_1_ and F_2_ hybrid fitness could easily be compared in a common environment (e.g., intermediate or ancestral) and the clear prediction is that the loss in fitness of F_2_ hybrids (due to segregating breakup of co-adapted compensatory alleles) compared to F_1_s will be greater in more divergently selected lines. One might also expect that selection for heterozygosity (for a given genomic hybrid index) will increase with parental divergence (Simon et al. 2018). Such an experiment would be valuable for establishing a general link between adaptive divergence and reproductive isolation.

Although I illustrate a correspondence between theory and data, I did so using a correlational approach and with a small sample size of 15 crosses. There are other plausible mechanisms besides pleiotropy that could underlie segregation variance in non-divergent characters. For example parallel phenotypic evolution (if it has a non-parallel genetic basis) can cause segregation variance in traits that do not differ between the parent taxa (Chevin et al. 2014; Thompson et al. 2019). For this mechanism to underlie the pattern shown in Fig. 2 there would have to be a correlation between parallel phenotypic evolution in some traits and divergent evolution in others — this seems unlikely. At the very least, my analysis should serve to buttress our assessment that models fundamentally based on pleiotropy such as Fisher’s (1930) geometric model are robust and useful abstractions of the evolutionary process.

## Acknowledgements

Feedback from D. Irwin, M. Osmond, S. Otto, J. Santangelo, D. Schluter, and S. Wang benefited the manuscript. K.Davis, N. Frasson, J. Heavyside, and M. Urquhart-Cronish assisted with the literature search (all) and/or data collection (M. U-C). I am grateful to all authors who responded to requests for data. R. Henriques created the bioR*χ*iv LATEXtemplate.

## Data accessibility

All data and analysis code used in this article will be deposited in a repository (e.g., Dryad) following publication. For now, they are available on my GitHub.

## Supplementary material 1: Supplementary methods

### A. Search strategy

I searched the literature for studies that made measurements of traits in F_1_ hybrids and their parents. To identify studies for possible inclusion, I conducted a systematic literature search using Web of Science. I included all papers that resulted from a general topic search of “Castle-Wright”, and from a topic search of “F_1_ OR hybrid OR inherit*” in articles published in *Evolution*, *Proceedings of the Royal Society B*, *Journal of Evolutionary Biology*, *Heredity*, or *Journal of Heredity*. These journals were selected because a preliminary search indicated that they contained nearly half of all suitable studies. These searches returned 106 studies deemed suitable after screening. To be more comprehensive, I conducted additional systematic searches by conducting similar topic searches among articles citing influential and highly-cited publications (Bradshaw et al. 1998; Churchill and Doerge 1994; Coyne and Orr 2004; Dobzhansky 1937; Grant 1981; Hatfield and Schluter 1999; Hubbs 1955; Lande 1981; Lynch and Walsh 1998; Mayr 1963; Schluter 2000; Tave 1986). The full literature search results are available in the archived data. My initial search returned 14048 studies, and after removing duplicates this left 11287 studies to be screened for possible inclusion. This literature search was primarily done for another unpublished study with the goal of understanding phenotype expression in F_1_ hybrids.

### B. Evaluation of studies

I required studies to meet several criteria to merit inclusion in my database. First, the study organisms had to originate recently from a natural (i.e., ‘wild’) population. This is because dominance patterns in domestic species differ substantially from non-domesticated species (Crnokrak and Roff 1995) and because I am explicitly interested in patterns as they occur in nature. I excluded studies using crops, domestic animals, laboratory populations that were > 10 (sexual) generations removed from the wild, or where populations were subject to artificial selection in the lab. If populations were maintained in a lab for more than 10 generations but were found by comparison to still strongly resemble the source population, I included the study (*n* = 2). I also excluded studies where the origin of the study populations was ambiguous. Hybrids had to be formed via the union of gametes from parental taxa, so I excluded studies using techniques like somatic fusion. Second, the ancestry of hybrids had to be clear. Many studies reported phenotypes of natural hybrids, for example in hybrid zones. I did not include these studies unless the hybrid category (i.e., F_1_, F_2_, backcross) was confidently determined with molecular markers (typically over 95 % probability, unless the authors themselves used a different cut-off in which case I went with their cut-off) or knowledge that hybrids were sterile and thus could not be beyond the F_1_).

Third, because I was interested in the inheritance of traits that are proximally related to organismal performance (McGee et al. 2015), I required studies to report measurements of at least one ‘non-fitness’ trait (’ordinary’ traits [Orr 2001]). Non-fitness traits (hereafter simply ‘traits’) are those that are likely under stabilizing selection at their optimum, whereas ‘fitness’ traits are those that are likely under directional selection and have no optimum (Merilä and Sheldon 1999; Schluter et al. 1991). In most cases it was possible to evaluate this distinction objectively because authors specifically referred to traits as components of fitness, reproductive isolating barriers, or as being affected by non-ecological hybrid incompatibilities. In some cases, however, I made the distinction myself. If particular trait values could be interpreted as resulting in universally low fitness, for example resistance to herbivores or pathogens, this trait was not included. The majority of cases were not difficult to assess, but I have included reasons for excluding particular studies or traits in the database screening notes (see Data accessibility).

Traits had to be measured in a quantitative manner to be included in the dataset. For example, if a trait was reported categorically (e.g., ‘parent-like; or ‘intermediate’), I did not include it. Some traits such as mate choice must often be scored discretely (in the absence of multiple trials per individual), even though the trait can vary on independent trials. Accordingly, we included discretely scored traits — like mate choice — when it was possible in principle to obtain a different outcome on independent trials. Such traits are recorded as 0s and 1s, but hybrids can be intermediate if both outcomes occurred with equal frequencies. I included traits where authors devised their own discrete scale for quantification. When suitable data were collected by the authors but not obtainable from the article, I wrote to the authors and requested the data. If the author cited a dissertation as containing the data, I attempted to locate the data therein because dissertations are not indexed by Web Of Science. I included multivariate trait summaries (e.g., PC axis scores) when reported. If traits reported both the raw trait values and the PC axis scores for a summary of those same traits, I collected both sets of data but omitted the PCs in our main analyses.

Using these criteria, I screened each article for suitability. As a first pass, I quickly assessed each article for suitability by reading the title and abstract and, if necessary, consulting the main text. After this initial search, I retained 407 studies. Since the previous steps were done by a team of five, I personally conducted an in-depth evaluation of each study flagged for possible inclusion. If deemed suitable, I next evaluated whether the necessary data could be obtained. After this second assessment, 198 studies remained. The reasons for exclusion of each study are documented the archived data (see Data accessibility).

### C. Data collection

For each study, I recorded several types of data. First, I recorded the mean, sample size, and an estimate of uncertainty (if available) for each measured trait for each parental crosses and hybrid category. In most cases, these data were included in tables or could be extracted from figures. In some cases, I contacted authors for the raw data or summary data. Each study conducted a minimum of three records to the larger database: one trait measured in each parent and the F_1_ generation. Traits were categorized as one of: behaviour, chemical, life history, morphological, physiological, or pigmentation. If the same traits were measured over ontogeny, I used only the final data point. When data were measured in multiple ‘trials’ or ‘sites’ I pooled them within and then across sites. If data were reported for different cross directions and/or sexes I recorded data for each cross direction / sex combination separately. Data processing was immeasurably aided by the functions implemented in the `tidyverse` (Wickham 2017).

For each paper I recorded whether the phenotypes were measured in the lab or field, if in the lab the number of generations of captivity, and whether a correlation matrix (preferably in recombinant – F_2_ or BC – hybrids; see below) was available or calculable from the raw data or figures. For the present study, specifically, each study would have had to contribute 8 or more datapoints – two traits from each of P_1_, P_2_, F_1_, and F_2_. Occasionally, different studies analysed different traits from individuals from the same crosses. In these cases, I simply grouped them as being the same study before analysis.

#### C.1. Comments on systematic nature of review

I attempted to follow PRISMA (Moher et al. 2009) guidelines to the best of my ability. Most of the criteria have been addressed above but a few other comments are warranted. I have no reason to suspect that any bias was introduced about estimates of parental divergence or segregation variance. This is because no studies seemed to have *a priori* hypotheses about such patterns. Accordingly, I do not believe that our estimates suffer from a file drawer problem, since detecting segregation variance in non-divergent traits was not the stated goal of any contributing studies. In addition, a formal meta-analytic framework here is not appropriate because I am not comparing studies that had any experimental treatment.

### D. Estimating genetic divergence and divergence time

I estimated genetic distance for pairs of species where data were available for both parents. A preliminary screening revealed that the internal transcribed spacer (ITS I and II) was the most commonly available gene for plants and cytochrome b was the most available gene for animals in our dataset. I downloaded sequences in R using the `rentrez` package (Winter 2017), and retained up to 40 sequences per species. Sequences were then aligned with the profile hidden Markov models implemented in the `align` function in the package, `aphid` (Wilkinson 2018). After aligning sequences I calculated genetic distance by simply counting the number of sites that differed between two aligned sequences, implemented using the the raw model option in the `dist.dna` function within `ape` (Paradis and Schliep 2018).

I also used timetree (Kumar et al. 2017) to obtain estimates of divergence time for each species pair in their database in years. After obtaining estimates of divergence time I regressed divergence time against the response and predictor variables used in the main analysis.

### E. Phylogenetic signal

To determine whether patterns might be spurious and caused by differences among taxa, I wished to see if there was phylogenetic signal in the data. If either my response or predictor variables co-varied with phylogeny this might indicate that phylogenetic independent contrasts or similar is necessary for analysis. We retrieved NCBI taxonomy IDs for our species using the `taxize R` package (Chamberlain and Szöcs 2013), and used these IDs (one arbitrarily chosen per cross) to generate a phylogeny using phyloT. Because branch lengths negligibly affect estimates of phylogenetic signal (Münkemüller et al. 2012), I assigned all branches equal lengths and used the `phylosig` function implemented in `phytools` (Revell 2012) to test for phylogenetic signal via Pagel’s *λ*.

## Supplementary material 2: Supplementary results

### A. Analyses with alternative analysis, filtering, and binning protocols

Although the analysis presented in the main text is, in my view, the most justifiable, there were several subjective decisions that I made when going from raw data to summary statistics. Accordingly, I wished to investigate the extent to which my findings were robust to making alternative choices. In this section I present a simple table showing the Spearman’s *ρ* coefficient and *P*-value for the correlation between phenotypic divergence in divergent traits and segregation variance in non-divergent traits. In all but one case, the pattern remains statistically significant.

There were correlations between sample size and some of my response variables, including estimates of parental divergence. To determine if my results were robust to the exclusion of potentially low-power studies, I generated two new datasets with studies measuring fewer than (i) 20 or (ii) 70 parental individuals excluded. Sample size did not predict parental divergence for these datasets. These results are presented below in Table S1 and indicate that sample size is not responsible for the pattern. I also note that a multiple regression with parental divergence and sample size as predictors — flawed because of potential heteroskedasticity and low data - predictor ratio — indicated that only parental divergence was a significant predictor of segregation variance (divergence *P* = 0.0195; sample size *P* = 0.755).

**Table S1.** Alternative data filtering and binning. All analyses with morphological traits only and binning based on statistical tests except where noted. The segregation variance of physiological traits and chemical traits was typically outside the magnitude observed in morphological traits, so we analyze the patterns for all traits and for all traits excluding physiological and chemical traits.

| Difference | Description | Spearman's $\rho$ | $P$ |
| --- | --- | --- | --- |
| Main text analysis | For reference | 0.8 | 0.0005809 |
| Less strict binning | Traits with $\geq 1$ SD difference between parents deemed 'divergent' | 0.714 | 0.00543 |
| Same segvar fomula; difference | Eqn. 1 used but difference not ratio | 0.729 | 0.00286 |
| Alternative segvar fomula; ratio | segvar = $\text{var}(F_2) / \text{var}(F_2)$ | 0.607 | 0.0186 |
| Alternative segvar fomula; difference | segvar = $\text{var}(F_2) - \text{var}(F_2)$ | 0.607 | 0.0186 |
| All traits included | No longer restricted to Morphology | 0.518 | 0.0523 |
| Most traits included | All traits except for physiology & life history (morphology, behaviour, pigment, life history) | 0.451 | 0.0364 |
| No transformation | No log transformation of variables | 0.746 | 0.00202 |
| Sample size filter #1 | No studies with fewer than 20 parent individuals | 0.736 | 0.00557 |
| Sample size filter #2 | No studies with fewer than 70 parent individuals | 0.762 | 0.0368 |

## Supplementary material 3: Supplementary figures

**Fig. S1.**
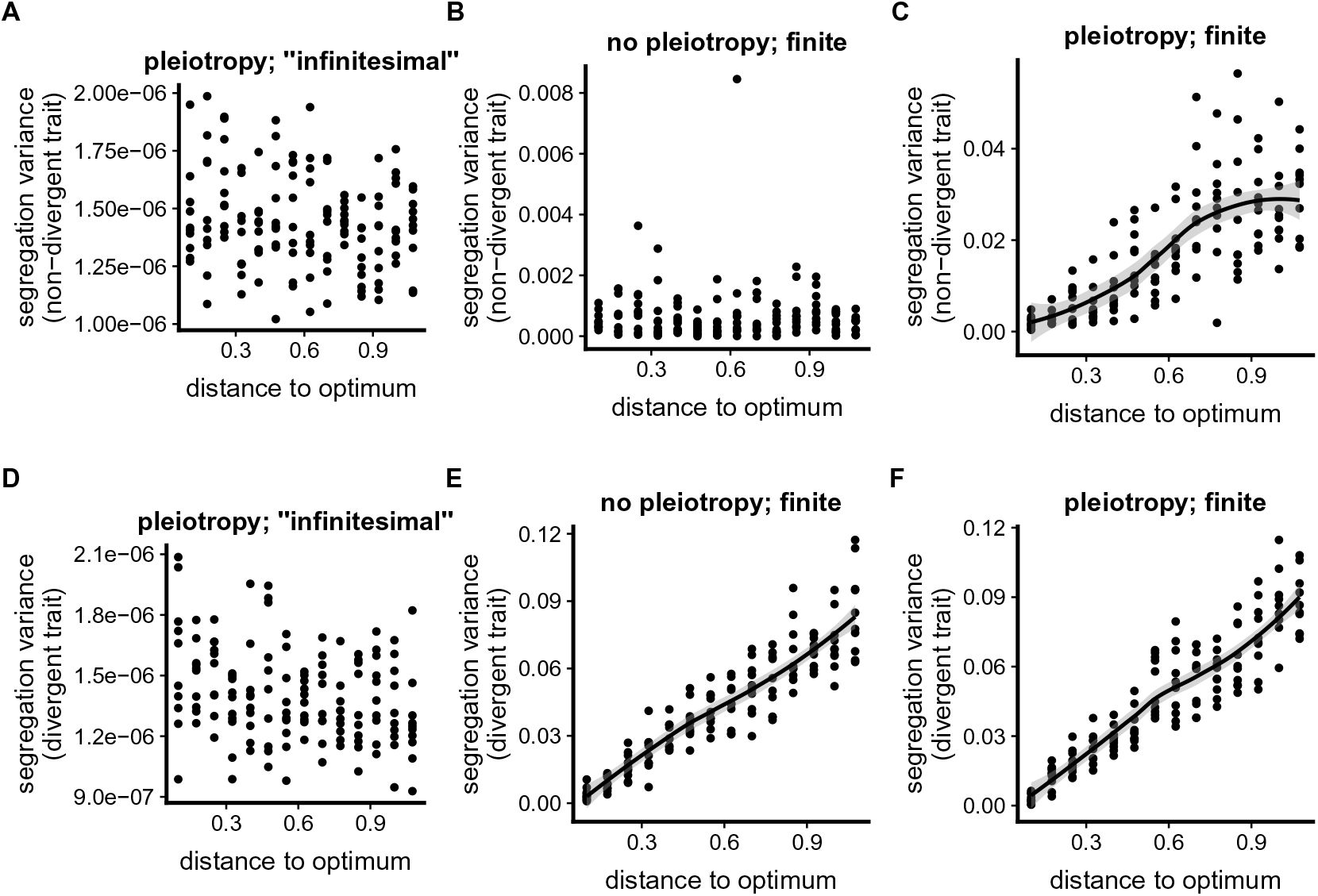
Simulation results to illustrate theoretical prediction. I conducted individual based simulations using methods similar to those described fully in Thompson et al. (2019), which is open access. Briefly, two initially identical populations diverged without gene flow between them for 1000 generations. All mutations were unique to each population (no parallelism). Individuals had two traits and only trait 1 was under selection. Optima were defined as [*d*, 0] for population 1 and [−*d*, 0] for population 2, where *d* is the distance to the optimum from the origin. I vary *d* along the horizontal axis in all figures. After 1000 generations I made interpopulation hybrids and measured the variance in traits 1 (*y*-axis, top row) and 2 (*y*-axis, bottom row). Panels **A** and **D** show simulations where populations adapt from pleiotropic but very small alleles. Panels **B** and **E** show the case where mutations are appreciably large but not pleiotropic— only affecting trait 1 or trait 2 but never both. Panels **C** and **F** show the case where mutations are appreciably large-effect and can affect both traits simultaneously. Simulation code is archived online. Simulations are haploid and so the F1 variance is the segregation variance. Here is what I would like you to get from the figure:(1) When mutations are small, there is no relationship between *d* and segregation variance in any trait, even with universal pleiotropy. (2) When mutations are large but there is no pleiotropy, a relationship between *d* and and segregation variance is observed only for the divergently-selected trait. And (3), with pleiotropic mutations of reasonably large effect, we see a relationship between *d* and and segregation variance for both traits. The only parameter that varies (other than the presence of pleiotropy) is mutation effect size; set to 10^−6^ in panels **A** and **D** and and 0.1 in the others.

**Fig. S2.**
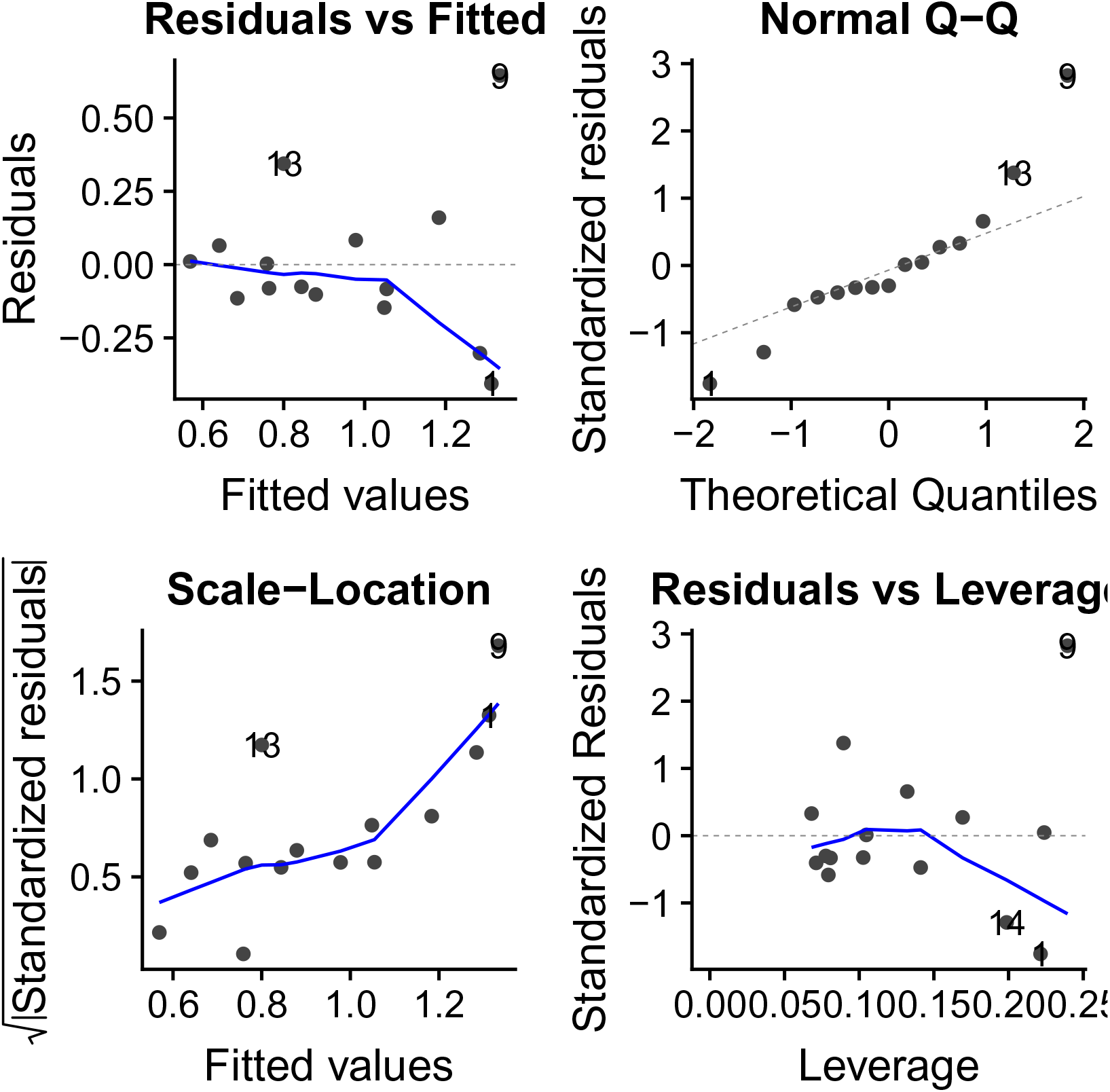
Diagnostics of the linear model testing the main hypothesis. The figure was produced with the `autoplot` function in the `ggfortify` R package. Although the linear regression is significant (F_1,12_ = 10.03, *r*^2^ = 0.455, *P* = 0.00812), the diagnostic plots reveal significant heteroskedasticity (Breusch-Pagan test *P* = 0.00182).

**Fig. S3.**
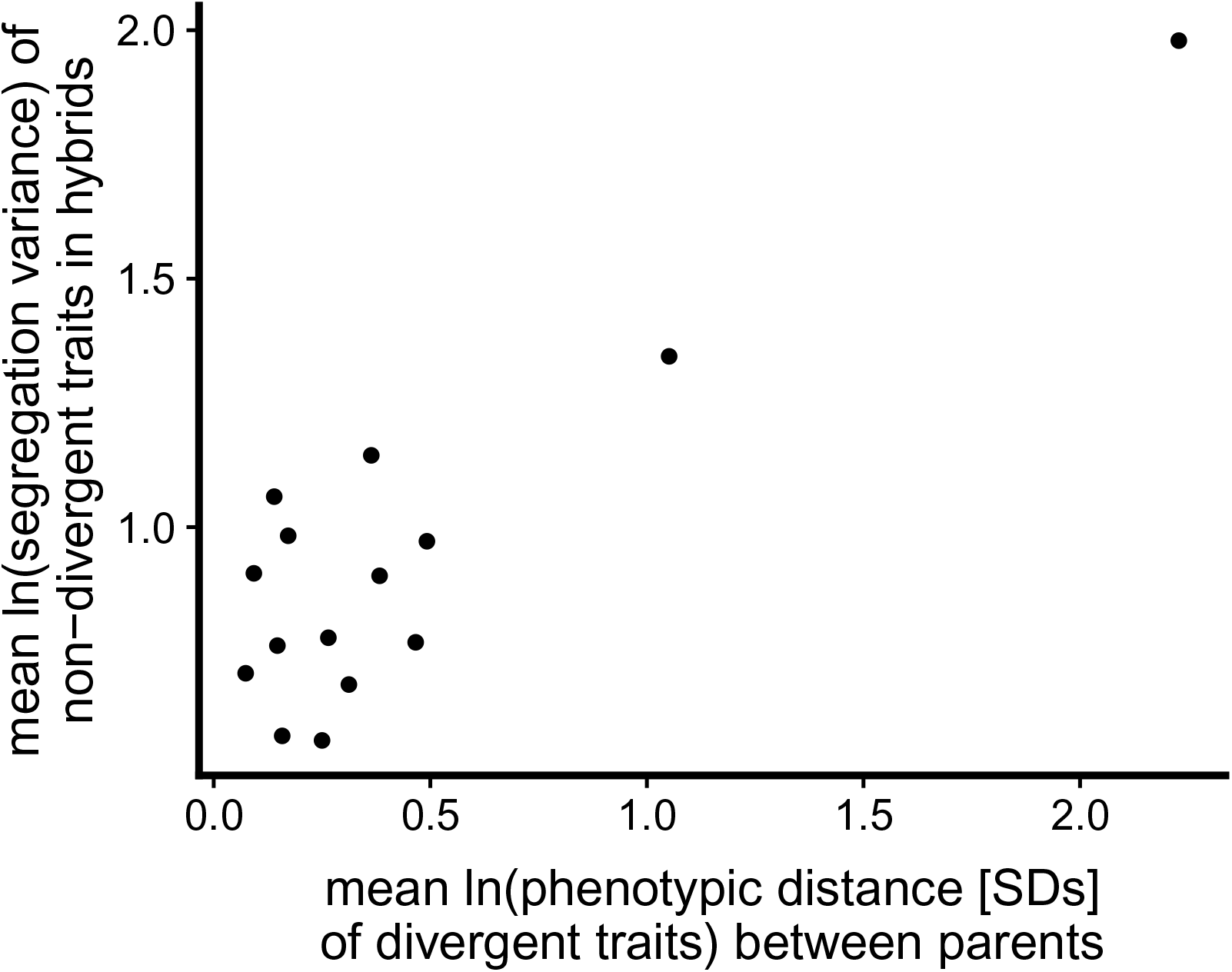
Segregation variance in non-divergent traits is not predicted by divergence in those traits. This analysis complements the analysis in the main text. A Spearman’s rank-order correlation is non significant (*ρ* = 0.446; *P* = 0.0972).

**Fig. S4.**
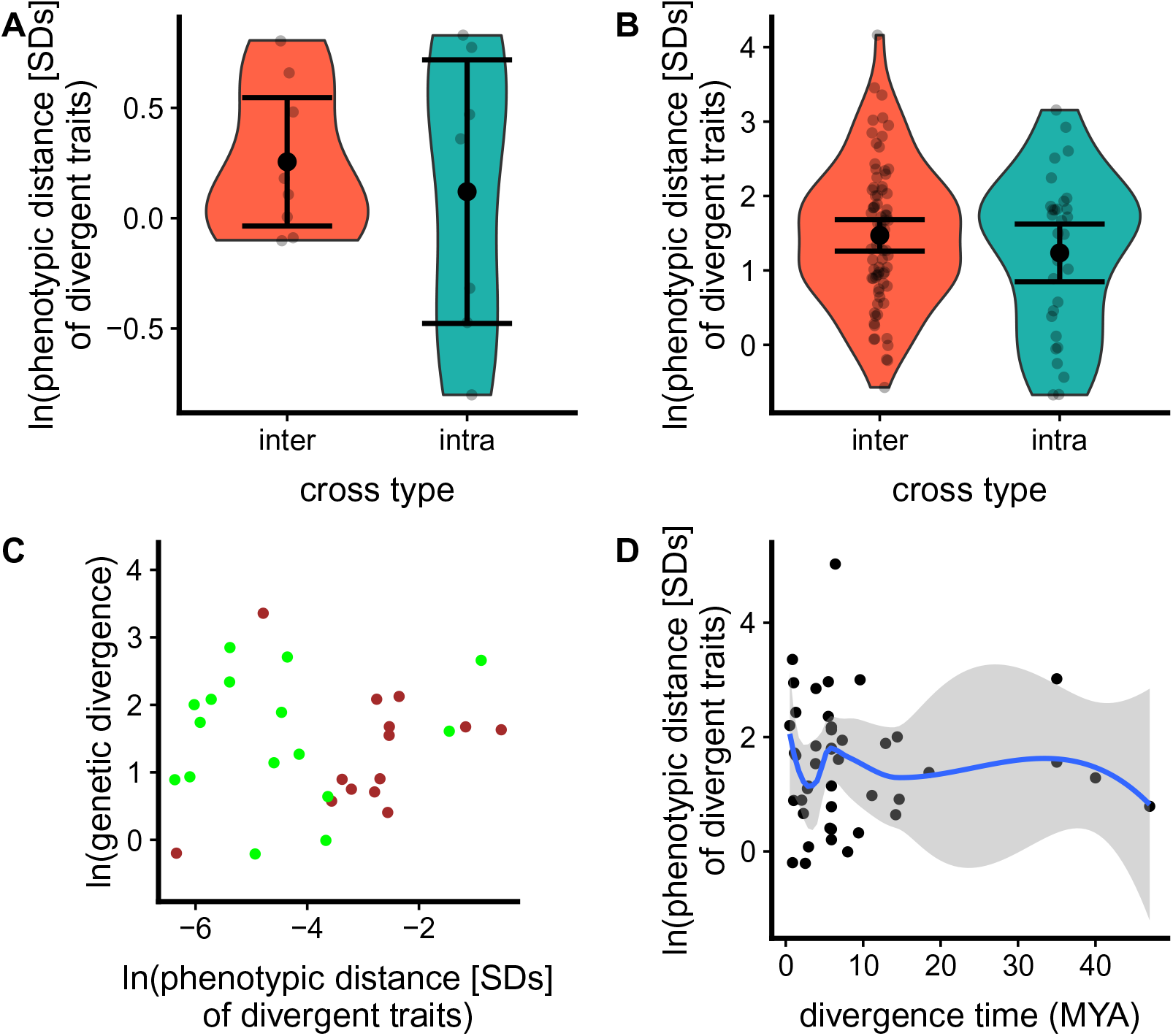
All available evidence suggests that phenotypic divergence between parents is uncorrelated with their divergence time. Panels **A** and **B** use intra- vs interspecific crosses as a proxy for divergence time. Panel **A** compares the taxa analyzed in the main text (*P* = 0.9…) and panel **B** uses a larger dataset of 198 studies (*P* = 0.268). Panel **C** uses continuous genetic distance between each pair of parents for which I could obtain DNA sequences in the larger database of 198 studies; sequence divergence does not predict parental phenotypic divergence (*P* = 0.746). Panel **D** uses estimates of divergence time from timetree for all pairs of species for which data were available; there is no relationship (*P* = 0.93). I calculated a unique value for each unique pair of species—if two studies crossed the same two species (regardless of subspecies or population status) the species pair only provides a single datum. For example, a study not included in the subset analyzed in the main text crossed *Drosophila simulans* to multiple *D. melanogaster* populations, but only provided a single datapoint for panels **B** and **C**.

